# An updated staging system for cephalochordate development: one table suits them all

**DOI:** 10.1101/2020.05.26.112193

**Authors:** João E. Carvalho, François Lahaye, Luok Wen Yong, Jenifer C. Croce, Hector Escrivá, Jr-Kai Yu, Michael Schubert

## Abstract

Chordates are divided into three subphyla: Vertebrata, Tunicata and Cephalochordata. Phylogenetically, the Cephalochordata, more commonly known as lancelets or amphioxus, constitute the sister group of Vertebrata and Tunicata. Lancelets are small, benthic, marine filter feeders, and their roughly three dozen described species are divided into three genera: *Branchiostoma*, *Epigonichthys* and *Asymmetron*. Due to their phylogenetic position and their stereotypical chordate morphology and genome architecture, lancelets are key models for understanding the evolutionary history of chordates. Lancelets have thus been studied by generations of scientists, with the first descriptions of adult anatomy and developmental morphology dating back to the 19^th^ century. Today, several different lancelet species are used as laboratory models, predominantly for developmental, molecular and genomic studies. Surprisingly, however, a universal staging system and an unambiguous nomenclature for developing lancelets have not yet been adopted by the scientific community.

In this work, we characterized the development of the European amphioxus (*Branchiostoma lanceolatum*) using confocal microscopy and compiled a streamlined developmental staging system, from fertilization through larval life, including an unambiguous stage nomenclature. By tracing growth curves of the European amphioxus reared at different temperatures, we were able to show that our staging system permitted an easy conversion of any developmental time into a specific stage name. Furthermore, comparisons of embryos and larvae from the European lancelet (*B. lanceolatum*), the Florida lancelet (*B. floridae*), the Chinese lancelet (*B. belcheri*), the Japanese lancelet (*B. japonicum*) and the Bahamas lancelet (*Asymmetron lucayanum*) demonstrated that our staging system could readily be applied to other lancelet species.

Although the detailed staging description was carried out on developing *B. lanceolatum*, the comparisons with other lancelet species thus strongly suggested that both staging and nomenclature are applicable to all extant lancelets. We conclude that this description of embryonic and larval development will be of great use for the scientific community and that it should be adopted as the new standard for defining and naming developing lancelets. More generally, we anticipate that this work will facilitate future studies comparing representatives from different chordate lineages.

## 1. Introduction

The subphylum Cephalochordata comprises only a few dozen species of small, lancet-shaped filter-feeders (Bertrand and Escrivá, 2011; Holland, 2015). The Cephalochordata (commonly referred to as lancelets or amphioxus) belong to the chordate phylum and are the sister group to all other chordates (Tunicata and Vertebrata) (Bertrand and Escrivá, 2011; Holland, 2015). Due to this phylogenetic position and their slow evolutionary rate (Louis et al., 2012), lancelets are considered valuable proxies for the chordate ancestor, both at the anatomic and genomic levels (Bertrand and Escrivá, 2011; Holland, 2015). The subphylum Cephalochordata is subdivided into three genera: *Branchiostoma*, *Epigonichthys* and *Asymmetron* (Poss and Boschung, 1996; Nishikawa, 2004; Zhang et al., 2006; Kon et al., 2007; Holland and Holland, 2010; Yue et al., 2014; Carvalho et al., 2017b; Subirana et al., 2020). Recent analyses of mitochondrial genomes suggested that the genus *Asymmetron* occupies the basal position and diverged from the *Epigonichthys / Branchiostoma* clade about 258-171 Mya (million years ago) (Subirana et al., 2020). It was further proposed that the split of the *Epigonichthys* and *Branchiostoma* lineages occurred about 182-120 Mya and that speciation within the genus *Branchiostoma*, between *B. belcheri* and *B. japonicum* versus *B. floridae* and *B. lanceolatum*, took place about 130-85 Mya (Subirana et al., 2020).

The importance of lancelets for understanding chordate evolution has driven generations of scientists to study their embryos and larvae (Holland and Holland, 2017). An initial description of lancelet development was already performed in the 19^th^ century, on *B. lanceolatum* material obtained in Naples, Italy (Kovalevsky, 1867). This work was subsequently completed, at the end of the 19^th^ and the beginning of the 20^th^ century, by a series of additional surveys on the same species (Hatschek, 1893; Cerfontaine, 1906; Conklin, 1932). More recently, in the early 1990s, the early development of *B. japonicum* was the subject of a detailed characterization by electron microscopy (Hirakow and Kajita, 1990, 1991, 1994). A similar approach was used to characterize neurulae, larvae and post-metamorphic specimens of *B. floridae* (Holland and Holland, 1992; Stokes and Holland, 1995). The most recent description of lancelet development was that of *A. lucayanum* embryos and larvae using differential interference contrast (DIC) microscopy (Holland and Holland, 2010; Holland et al., 2015). Taken together, these studies on species of the two most distantly related lancelet genera have revealed that the ontogeny of lancelets is a highly coordinated and conserved process. It is thus all the more surprising that there is currently no universal developmental staging system available for the members of this subphylum.

In the course of the last three decades, lancelets have become important models for addressing developmental processes from a molecular and genomic perspective (Bertrand and Escrivá, 2011; Acemel et al., 2016; Carvalho et al., 2017b; Marlétaz et al., 2018; Simakov et al., 2020). However, unlike for other developmental model organisms, such as zebrafish, the scientific community is using different lancelet species for their studies, with the choice being mainly dependent on the availability of animal resources (Carvalho et al., 2017b). Husbandry protocols have been established for at least five lancelet species (Carvalho et al., 2017b), but, due to the absence of a universal staging system, the nomenclature of embryos and larvae obtained with these protocols has become extremely confusing. While developing lancelets are often named in accordance with previous reports on the same species (Bertrand et al., 2011; Lu et al., 2012; Holland, 2015; Annona et al., 2017), it is also not uncommon to indicate the time after fertilization, usually measured in hours after fertilization (Fuentes et al., 2007; Bertrand and Escrivá, 2011). However, developmental speed is known to vary between lancelet species and to depend on the rearing temperature, which is not the same in each study (Fuentes et al., 2007; Bertrand and Escrivá, 2011). The absence of an unambiguous nomenclature for developing lancelets artificially complicates comparisons of results obtained in different species and sometimes even within the same species, for example, when two laboratories use incompatible staging styles (Bertrand et al., 2011; Pantzartzi et al., 2017). There is, therefore, an urgent need to establish an easy and systematic classification for embryonic and larval development that applies to different lancelet species.

To achieve this objective, we illustrated the development of *B. lanceolatum* using confocal microscopy and established growth curves at different temperatures based on the number of somites. We further compared embryos and larvae of *B. lanceolatum* with those of other lancelets. By applying and expanding the stage definitions of Hirakow and Kajita (Hirakow and Kajita, 1990, 1991, 1994) and Lu and colleagues (Lu et al., 2012), we compiled a streamlined staging system of *B. lanceolatum* development, from fertilization through larval life, with an unambiguous stage nomenclature. Analyses of the growth curves revealed that our staging system could be used to easily convert developmental times into unambiguous stage names, at any given rearing temperature. Furthermore, comparisons between *B. lanceolatum*, *B. floridae*, *B. belcheri*, *B. japonicum* and *A. lucayanum* embryos and larvae demonstrated that the updated staging system could readily be applied to other lancelet species. We hope that the scientific community will adopt this universal developmental staging system for lancelets to facilitate the use of these fascinating animals as laboratory models.

## 2. Material and methods

### 2.1. Animal husbandry and *in vitro* cultures

Ripe *B. lanceolatum* adults were collected by dredging in Argelès-sur-Mer, France, and retrieved from the sand by sieving. Animals were transported, quarantined and maintained in Villefranche-sur-Mer as previously described (Carvalho et al., 2017b). Spawning was induced by a 36-hour thermal shock at 23°C (Fuentes et al., 2004). Sperm and oocytes were collected separately and fertilization was performed *in vitro*. *B. lanceolatum* embryos and larvae were raised in the dark at constant temperatures (16°C, 19°C or 22°C) until the desired developmental stages, and larvae were fed daily with *Tisochrysis lutea* algae (Carvalho et al., 2017b).

Adult *B. floridae* were collected in Tampa Bay, Florida, USA. Animals were maintained in the laboratory as previously described (Zhang et al., 2007; Yong et al., 2019). Gametes were obtained either by electric stimulation, heat shock or spontaneous spawning (Holland and Yu, 2004; Ono et al., 2018). Embryos and larvae were cultured at constant temperatures (25°C or 30°C) until the desired stages, and larvae were fed daily with *Isochrysis sp.* algae.

Adult *B. belcheri* and *B. japonicum* were collected in Kinmen Island near Xiamen in southeastern China (Zhang et al., 2013). Animals were maintained as previously described (Zhang et al., 2007; Yong et al., 2019). Embryos were obtained through spontaneous spawning in the facility (Zhang et al., 2007). Embryos and larvae were cultured at a constant temperature (24°C for *B. belcheri* and 25°C for *B. japonicum*) until the desired stages, and larvae were fed daily with *Isochrysis sp.* algae.

*A. lucayanum* adults were collected in the lagoon between North and South Bimini, Bahamas. Embryos and larvae were obtained and subsequently cultured at a constant temperature (27°C) as previously described (Holland and Holland, 2010).

### 2.2. Differential interference contrast (DIC) microscopy

Embryos and larvae used for observation and imaging by DIC microscopy were fixed in 4% PFA in MOPS buffer for 1 hour at room temperature or overnight at 4°C. Embryos and larvae were subsequently washed twice in ice-cold 70% ethanol in DEPC-treated water and stored at −20°C until further use. Embryos and larvae were rehydrated in PBS buffer and mounted in PBS buffer or 80% glycerol for imaging.

DIC microscopy of *B. lanceolatum* embryos and larvae was performed using a Zeiss Axiophot microscope, equipped with an AxioCam ERc 5s camera (Carl Zeiss SAS, Marly-le-Roi, France). Images of *B. floridae*, *B. belcheri*, *B. japonicum* and *A. lucayanum* embryos and larvae were acquired with a Zeiss Axio Imager A1 microscope, equipped with a AxioCam HRc CCD camera (Carl Zeiss SAS, Marly-le-Roi, France). For 64-cell, 128-cell and blastula stages, multiple z-levels were taken manually. The z-stack images were processed with the Extended-Depth-of-Field plugin of the ImageJ software using default settings (Schneider et al., 2012), and panels were subsequently formatted with Adobe Photoshop CS6 (Adobe Inc., San Jose, USA).

### 2.3. Fluorescent staining and immunohistochemistry

*B. lanceolatum* fertilized egg, cleavage- and gastrula-stage embryos were stained using FM 4-64 lipophilic dye (Invitrogen, Cergy Pontoise, France) at a final concentration of 10 μg/ml. The FM 4-64 lipophilic dye is a nontoxic vital dye commonly used to label plasma membranes and endocytic pathways (Sardet et al., 2011). Following dye incubation, the embryos were fixed for 1 hour at room temperature with freshly prepared 4% PFA (paraformaldehyde) in MOPS buffer (Yu and Holland, 2009). Embryos were washed twice in 70% ethanol and subsequently rehydrated in PBS buffer (Yu and Holland, 2009). Nuclear DNA staining was performed for 10 minutes at room temperature using Hoechst dye (Invitrogen, Cergy Pontoise, France) at a final dilution of 1:5000. Embryos were mounted in PBS buffer and imaged within 3 hours after staining with the FM 4-64 and Hoechst dyes.

For neurula, tailbud and larva stages, the FM 4-64 lipophilic dye yielded unsatisfactory results. These stages were thus stained by immunohistochemistry using a primary antibody against aPKC (polarity protein atypical protein kinase C), which labels structures associated with cell membranes (Patalano et al., 2006; Prulière et al., 2011). For whole-mount immunohistochemistry *B. lanceolatum* embryos and larvae were fixed overnight at 4°C in freshly prepared ice-cold 4% PFA in MOPS buffer (Yu and Holland, 2009). Immunohistochemistry was performed as previously described (Zieger et al., 2018), using the primary antibody against aPKC (SC216, Santa Cruz Biotechnology, Dallas, USA) at a final dilution of 1:100 and a secondary anti-mouse IgG-heavy and light chain antibody conjugated with Cy3™ (A90-516C3, Bethyl Laboratories Inc., Montgomery, USA) at a final dilution of 1:200. Hoechst dye (Invitrogen, Cergy Pontoise, France) at a final dilution of 1:5000 was used for nuclear DNA staining. Embryos were mounted in PBS buffer and subsequently imaged.

Imaging was systematically carried out on a Leica TCS SP8 confocal microscope, using a 20x objective (0.75 IMM HC PL APO CORR CS WD = 0,68mm) (Leica Microsystems SAS, Nanterre, France). FM 4-64/DNA staining and aPKC/DNA staining scans were obtained sequentially. DNA, FM 4-64 and aPKC staining were excited using, respectively, 405nm, 514nm and 552nm lasers. Series of optical sections were taken at a z-step interval of 2 μm. The ImageJ software (Schneider et al., 2012) was subsequently used for image processing and to generate maximum as well as average projections. Adobe Photoshop CS6 (Adobe Inc., San Jose, USA) was used to format larger panels requiring the reconstitution of partial images.

### 2.4. Growth curves and *in situ* hybridization

Developing *B. lanceolatum* embryos were reared at three different temperatures: 16°C, 19°C and 22°C. At regular intervals, animals were collected and fixed for subsequent *in situ* hybridization analyses. A 874-bp fragment containing the complete coding sequence of the *B. lanceolatum mrf1* (*myogenic regulatory factor 1*) gene, a member of the *myoD* gene family (Schubert et al., 2003), was amplified by PCR from cDNA and cloned into the pGEM-T Easy Vector (GenBank accession number of *B. lanceolatum mrf1*: MT452570). *In situ* hybridization experiments were carried out with a *mrf1*-specific antisense riboprobe as previously described (Yu and Holland, 2009; Carvalho et al., 2017c). Following *in situ* hybridization, embryos and larvae were mounted for DIC microscopy and imaged as described above.

Expression of the *mrf1* gene was used to visualize the somites and thus to obtain somite pair counts of embryos and larvae reared at different temperatures. The somite pair counts were used to define a training set of data points for each rearing temperature (16°C, 19°C and 22°C), hence allowing the calculation of best natural logarithmic tendency curves using Microsoft Excel (Microsoft Corporation, Redmond, USA). The curves were subsequently curated and used to define time intervals for each developmental stage (i.e. cleavage, blastula, gastrula, neurula, tailbud and larva stages).

## 3. Results

### 3.1. *Branchiostoma lanceolatum* staging series

Making use of the available *in vitro* culture protocols for developing lancelets (Carvalho et al., 2017b), the updated staging system was established using *B. lanceolatum* embryos and larvae. Prior to confocal imaging, embryos and larvae were fixed at the desired stages and labeled with fluorescent probes marking cell membranes and nuclei, hence allowing detailed morphological analyses of individual developmental stages. In the following, each stage of the updated staging system will be presented and defined. The stage names are indicative of the developmental period and are in accordance with previous descriptions of lancelet development (Hirakow and Kajita, 1990, 1991, 1994; Lu et al., 2012) as well as with the recently developed ontology for the *Branchiostoma* genus, AMPHX (Bertrand et al., 2021).

#### 3.1.1. Fertilization and cleavage

Lancelets are gonochoric and reproduce by external fertilization. Under appropriate environmental conditions, gravid males and females respectively release mature spermatozoa and oocytes into the water column. Prior to spawning, the mature lancelet oocyte undergoes the first meiotic division with formation of the first polar body and it is subsequently arrested in the second meiotic metaphase (Holland and Onai, 2012). Following spawning, the second meiotic division of the oocyte is triggered by fertilization and is completed within 10 min. The second meiotic division leads to the formation of the second polar body and the migration of the maternal chromosomes to the animal pole, which is defined by the position of the polar body (Fig. 1A, Supplementary Fig. 1A) (Holland and Holland, 1992). Independent of the entry point, the nucleus of the sperm first migrates to the vegetal half and only then joins the maternal chromosomes at the animal pole (Holland, 2015). Very soon after fertilization, a whorl composed of sheets of endoplasmic reticulum is further formed within the 1-cell stage. This whorl likely constitutes the germ plasm, since expression of germ cell markers, such as *nanos* and *vasa*, is associated with this structure (Wu et al., 2011). The 1-cell stage embryo is semi-opaque, due to the high quantity of granules uniformly distributed throughout the cell, and is surrounded by a membrane called the vitelline layer (Willey, 1894). As soon as fertilization occurs, the vitelline layer detaches from the 1-cell stage and expands, giving rise to the fertilization envelope (Holland and Holland, 1989). Cleavage, gastrulation and the first stages of neurulation will occur within the fertilization envelope (Holland, 2015).

**Figure 1.**
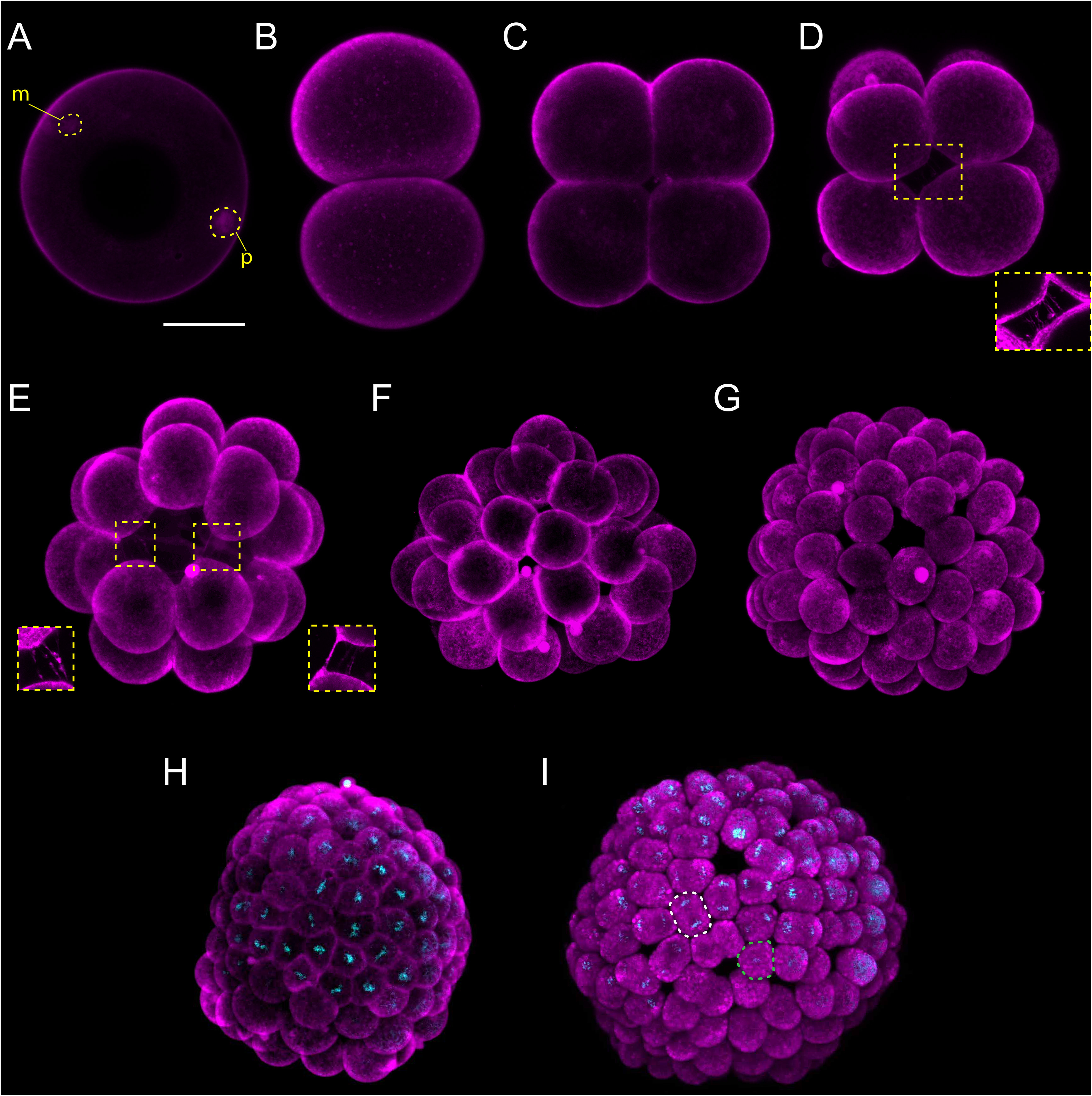
*Branchiostoma lanceolatum* fertilization, cleavage and blastula stages. Embryos are stained with the lipophilic dye FM 4-64 (magenta). (B,C) Animal pole views. (D-I) Animal pole is up. Maximum projections of confocal z-stacks of *B. lanceolatum* embryos at the (A) 1 cell-stage, (B) 2-cell stage, (C) 4-cell stage, (D) 8-cell stage, (E) 16-cell stage, (F) 32-cell stage, (G) 64-cell stage, (H) 128-cell stage and (I) blastula stage. Insets in (D) and (E) show slender filopodia between blastomeres. In (H,I), Hoechst DNA staining (cyan) shows synchronous cell divisions at the 128-cell stage (H) and asynchronous cell divisions at the forming blastula B-stage (I), with a white dashed line highlighting a cell in telophase and a green dashed line highlighting a cell following cytokinesis. Abbreviations: m – maternal DNA; p – paternal DNA. Scale bar: 50 μm.

Lancelet cleavage is radial holoblastic, meaning that cleavage completely separates blastomeres and results in early stage embryos with radial symmetry along the animal-vegetal axis (Barresi and Gilbert, 2019). The first cleavage starts from the animal pole and gives rise to the 2-cell stage, which is composed of two identically shaped blastomeres (Fig. 1B). When dissociated, each one of the first two blastomeres can give rise to a complete animal, but only one of the two blastomeres inherits the germ plasm (Holland and Onai, 2012). The second division is meridional and at a right angle to the first one, creating four blastomeres with approximately equal size, the 4-cell stage (Fig. 1C). Individual blastomeres are not adhering very strongly at this stage, and their dissociation can lead to the formation of twins or even quadruplets (Holland and Onai, 2012). Cleavage continues by an equatorial division, creating four animal and four vegetal blastomeres at the 8-cell stage, with the former being smaller than the latter (Fig. 1D). The blastomeres are held together by short microvilli and slender filopodial processes that bridge the space between adjacent blastomeres (insets in Fig. 1D,E) (Hirakow and Kajita, 1990). The 16-cell stage is the result of a meridional cleavage (Fig. 1E), and the 32-cell stage of a subsequent equatorial cleavage of each blastomere (Fig. 1F). At the 32-cell stage, the embryo is composed of a single layer of cells forming a central cavity called the blastocoel (Supplementary Fig. 1B) (Grassé, 1948; Hirakow and Kajita, 1990). The blastomeres will keep dividing regularly, giving rise to the 64-cell stage (Fig. 1G) and then to the 128-cell stage (Fig. 1H). The 8^th^ cell division cycle, i.e. the transition from 128 cells to 256 cells, which we define as the B stage, is characterized by the initiation of asynchronous cell division within the embryo (Grassé, 1948; Hirakow and Kajita, 1990) and further marks the formation of the blastula (Fig. 1I). The cells constituting the blastula will divide further, until the initiation of the gastrulation process.

#### 3.1.2. Gastrulation

The cells forming the hollow blastula are not identical in shape and size. The vegetal blastula cells are larger and hence indicate where the initial flattening of the gastrula takes place at the G0 stage (Fig. 2A) (Willey, 1894; Holland, 2015). The vegetal side of the embryo will continue to flatten and bend inward at the G1 stage (Fig. 2B,B’), hence forming a depression that marks the position of the blastopore. Thereafter, the vegetal tissue starts to invaginate into the blastocoel at the G2 stage (Fig. 2C,C’) (Hirakow and Kajita, 1991). The invaginating cells correspond to the presumptive endomesoderm, while the non-invaginating cells of the outer layer constitute the future general and neural ectoderm (Holland and Onai, 2012). As gastrulation proceeds with further cell divisions, the invaginating cells reduce the size of the blastocoelic cavity, ultimately leading, at the G3 stage, to a two-layered gastrula with an archenteron and a blastoporal lip. In this cap-shaped gastrula, the diameter of the blastopore is about half the size of the entire embryo (Fig. 2D,D’) (Hirakow and Kajita, 1991). Subsequent gastrulation movements result in an expansion of the cavity of the archenteron and in an almost complete loss of the blastocoelic cavity. This process leads to a narrowing of the blastoporal opening, which inflects the blastoporal lip, forming a cup-shaped gastrula at the G4 stage (Fig. 2E,E’) and a vase-shaped gastrula at the G5 stage (Fig. 2F,F’) (Hirakow and Kajita, 1991). Starting at the G5 stage, differences between the dorsal and ventral sides of the embryo become discernable, with the dorsal side beginning to flatten (Fig. 2F,F’) (Willey, 1894). These differences become more pronounced at the G6 stage, as the size of the blastopore continues to decrease and the embryo continues to elongate (Fig. 2G,G’). At this late gastrula stage, the embryo is bottle-shaped, and the blastopore starts to incline towards the dorsal side of the embryo, which is likely a synapomorphic trait of chordates, already present in their last common ancestor (Willey, 1894).

**Figure 2.**
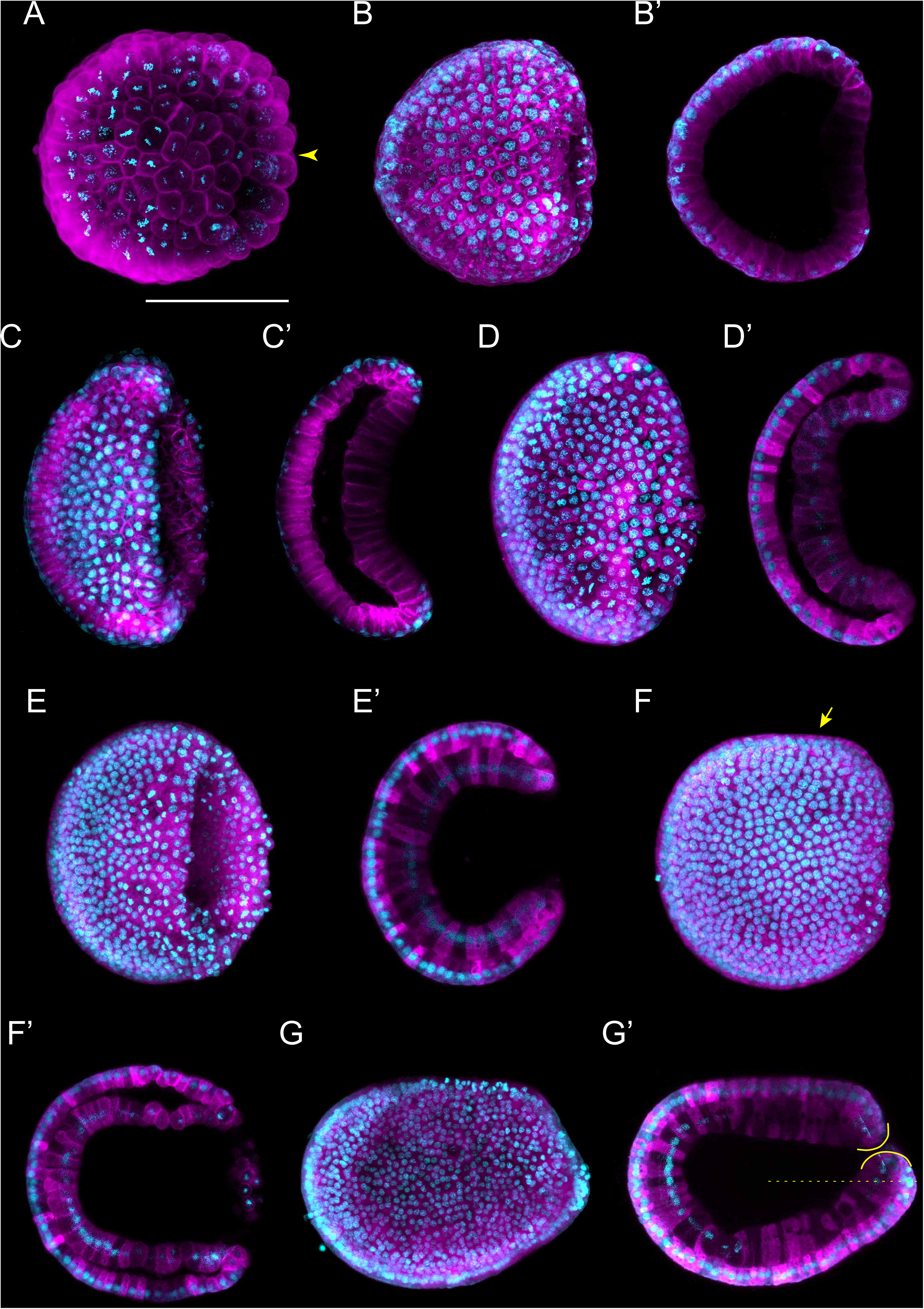
*Branchiostoma lanceolatum* gastrula stages. Embryos are stained with the lipophilic dye FM 4-64 (magenta) and with the DNA dye Hoechst (cyan). Animal pole and anterior pole are to the left and dorsal side is up. (A-G) Maximum projections of confocal z-stacks of entire embryos (B’-G’). Single z-stacks highlighting the inner morphology of the developing gastrula. (A) G0 stage, (B,B’) G1 stage, (C,C’) G2 stage, (D,D’) G3 stage, (E,E’) G4 stage, (F,F’) G5 stage, (G,G’) G6 stage. In (A), the yellow arrowhead indicates the vegetal cells. In (F), the yellow arrow highlights the flattened side of the gastrula embryo. In (G’), the yellow lines delimit the upper and lower lips of the blastopore, and the dashed line indicates the midline of the embryo. Scale bar: 50 μm.

Expression patterns of marker genes have determined that, with the exception of the tissues located in the immediate vicinity of the blastopore, most of the gastrula is destined to become the anteriormost region of the amphioxus larva. This includes the lancelet cerebral vesicle, the anteriormost somites, the pharynx with mouth and gill slits as well as the anterior section of the notochord (Holland and Onai, 2012). Transplantation experiments further demonstrated that the dorsal lip of the blastopore corresponds to a gastrulation organizer, similar or equivalent to the Spemann-Mangold organizer of vertebrates (Tung et al., 1961, 1962; Le Petillon et al., 2017).

#### 3.1.3. Neurulation

Following gastrulation, ectodermal cells develop cilia (Supplementary Fig. 1C,C’), and the embryo therefore starts to rotate within the fertilization envelope by ciliary movement (Lu et al., 2012; Holland, 2015). Cilia are also present on the endomesodermal cells of the archenteron (Hirakow and Kajita, 1991), and these cilia have been shown to play a role in establishing left-right asymmetry (Blum et al., 2014; Zhu et al., 2020). At this point in development, the N0 stage, neurulation starts. The N0 stage embryo is unsegmented and shows a typical dipoblastic organization, with the ectoderm externally and the endomesoderm internally (Fig. 3A). A small blastopore is still visible, and the dorsal ectoderm, destined to become the neuroectoderm, is flat with a shallow longitudinal groove (Fig. 3A). The subsequent N1 stage is characterized by the establishment of the first somites (somite pairs 1 through 3) (Fig. 3B,B’). The mesoderm, located dorsally within the endomesoderm, forms three folds: one medially that will develop into the notochord and two laterally that will give rise to the anterior somite pairs (Supplementary Fig. 1C’). At the N1 stage, the somites start pinching off in an anterior to posterior sequence. At the same stage, the dorsal non-neural ectoderm starts to detach from the edges of the neural plate. Following their detachment, the ectodermal cells will migrate over the neural plate using lamellipodia and fuse at the dorsal midline (Holland et al., 1996). At the end of this process, the neural plate will be completely covered by non-neural ectoderm, and the neuropore will have been formed anteriorly (Supplementary Fig. 1C) (Hatschek, 1881, 1893; Holland and Onai, 2012).

**Figure 3.**
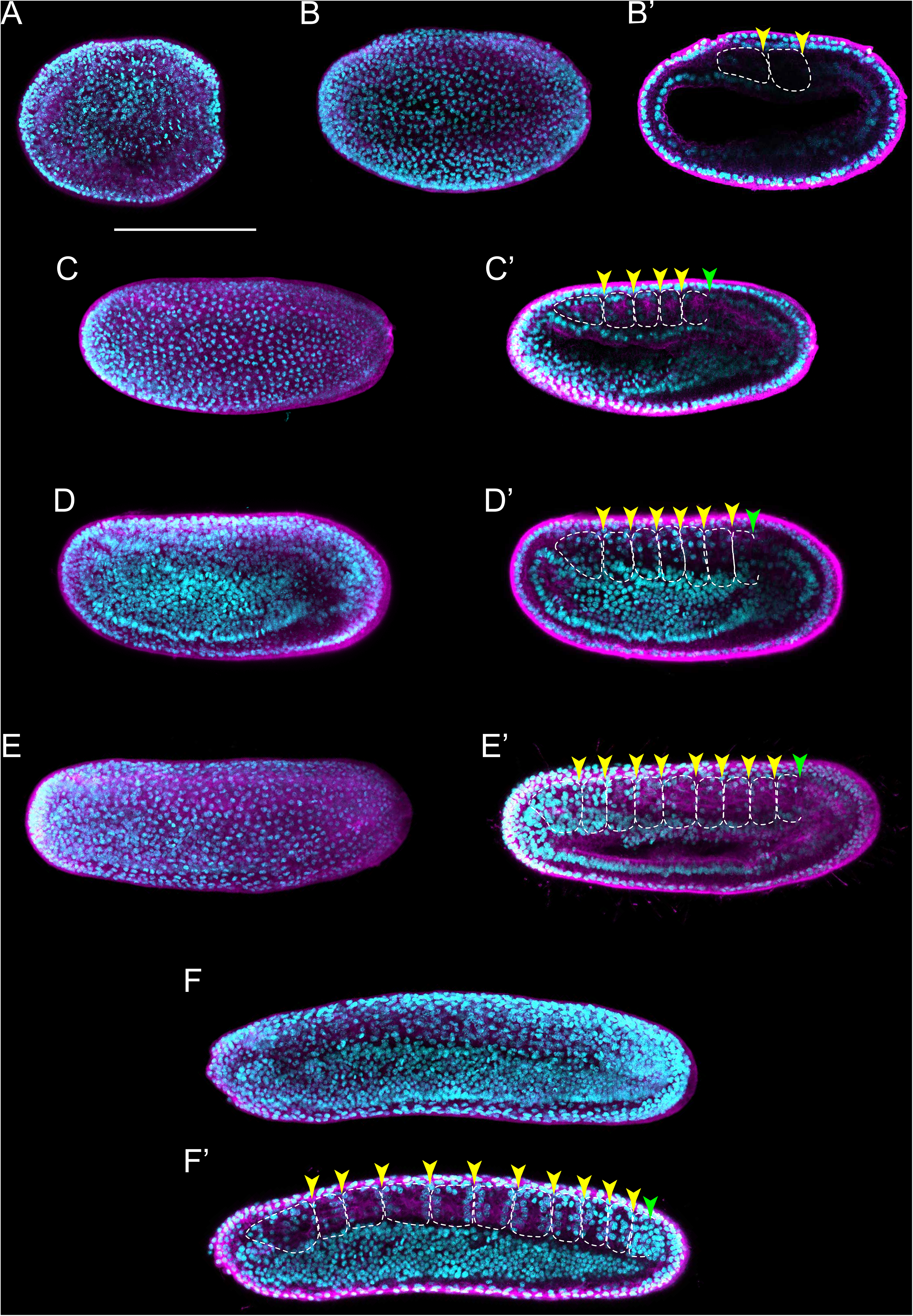
*Branchiostoma lanceolatum* neurula stages. Embryos are labeled for aPKC (magenta) and stained with the DNA dye Hoechst (cyan). Anterior pole is to the left and dorsal side is up. (A-G) Average projections for aPKC (magenta) and maximum projections for Hoechst DNA staining (cyan) of confocal z-stacks of entire embryos. (B’-G’) Single z-stacks highlighting the inner morphology of the developing neurula. (A) N0 stage, (B,B’) N1 stage, (C,C’) N2 stage, (D,D’) N3 stage, (E,E’) N4 stage, (F,F’) N5 stage. In (B’-F’), white dashed lines delineate the somites, the yellow arrowheads indicate the posterior limit of the somites and the green arrowheads highlight the posterior limit of somites newly established by enterocoely (C’-E’) or newly formed by schizocoely (F’). Scale bar: 100 μm.

As neurulation proceeds, the archenteron is no longer in contact with the exterior, but still communicates with the forming neural tube: the blastopore is incorporated into the neurenteric canal, which connects the neural tube with the archenteron (Supplementary Fig. 1D,E,E’,E’’), which becomes the presumptive gastric cavity (Willey, 1894). The embryo keeps elongating by the addition of new somites, reaching 4 to 5 somite pairs at the N2 stage (Fig. 3C,C’, Supplementary Fig. 1E’). At this stage, the embryo hatches from the fertilization envelope by the synthesis and secretion of hatching enzymes and starts swimming freely by ciliary activity (Stokes and Holland, 1995; Stokes, 1997). The neural plate is V-shaped (Supplementary Fig. 1E) and the primordium of the notochord is a round mass of cells extending ventrally along the neural plate (Supplementary Fig. 1E’). Central nervous system, notochord and somites are clearly distinguishable, although the boundaries between notochord and somites are not always evident (Fig. 3C’, Supplementary Fig. 1E,E’,E’’) (Hirakow and Kajita, 1994). The archenteron located anterior to the first somite pair starts expanding at this stage, forming two dorsolateral lobes (Supplementary Fig. 1E’’).

At the N3 stage, the embryo is characterized by 6 to 7 somite pairs (Fig. 3D,D’). The neural tube is closing, but will only become circular at subsequent developmental stages. The notochord is individualized from the somites, except at the most anterior tip of the embryo (Hatschek, 1893; Conklin, 1932). Ventral extensions of the somites start to generate the lateral and ventral coeloms as well as the musculature of the atrial floor (Holland and Onai, 2012). Furthermore, expression of early markers of Hatschek’s nephridium, such as *pax2/5/8*, becomes detectable in the mesothelial wall of the first somite on the left side of the embryo (Kozmik et al., 1999, 2007; Carvalho et al., 2017a). The subsequent N4 stage is characterized by 8 to 9 somite pairs (Fig. 3E,E’). At this stage, the two dorsolateral lobes that originated from the anterior archenteron have formed two distinctive head cavities: Hatschek’s left and right diverticulum (Willey, 1894; Grassé, 1948).

The N5 stage, which is characterized by 10 to 11 somite pairs, is when the asymmetric formation of somites from the tail bud is initiated (Fig. 3F,F’). Thus, while early somites are established from endomesoderm internalized during gastrulation by enterocoely, starting at the N5 stage, somites are formed by schizocoely from the tail bud (Holland, 2015). At this stage, the left and right diverticulum are asymmetrically organized: while the left diverticulum roughly maintains its original form and size, the right diverticulum moves anteriorly, flattens and increases in size (Willey, 1894). Furthermore, the primordium of the club shaped gland is first discernable, ventrally in the anterior endoderm on the right side of the embryo. This developmental stage is further characterized by a decrease of proliferative activity in somites and notochord, where it becomes limited to cells at the posterior end of the embryo. However, cell proliferation continues in the tail bud, in the endoderm and in the anterior neural plate (Holland and Holland, 2006).

#### 3.1.4. Tailbud and larva

Following neurulation, at the T0 stage, the embryo has 12 pairs of somites and exhibits a transitional morphology between neurula and larva stages (Fig. 4A,A’) that resembles a generic vertebrate tailbud stage embryo (Slack et al., 1993; Marlétaz et al., 2018). At this T0 stage, the anterior portion of the embryo becomes clearly distinct from the posterior one, as the pharyngeal region commences to grow. In addition, the embryo starts to twitch and bend as its neuromuscular system slowly becomes operational (Hirakow and Kajita, 1994). At the subsequent T1 stage, embryos are longer than those at the T0 stage, but this length difference is not due to the addition of a significant number of new somite pairs. Instead, it is due to the maturation and elongation of the existing ones, in particular those located in the anterior half of the embryo (Fig. 4B,B’, Supplementary Fig. 2A,B). The overall shape of the embryo also changes at the T1 stage: the body is becoming slender as the embryo elongates, a distinctive rostral snout is appearing and the tail fin is starting to form in the caudal ectoderm (Supplementary Fig. 2B) (Hirakow and Kajita, 1994). The first pigment spot in the central nervous system appears, located in the ventral wall of the neural tube at the level of the fifth somite pair (Supplementary Fig. 2B) (Willey, 1894). Concomitant with the elongation of the rostral snout, the right diverticulum expands anteriorly, hence forming the snout cavity below the notochord (Supplementary Fig. 2A,C). In addition, the left diverticulum starts fusing with the ectoderm to form the pre-oral pit, and the anlage of the mouth is clearly visible. Yet, neither one of these two structures penetrates the ectoderm and opens to the exterior at this stage (Kaji et al., 2016).

**Figure 4.**
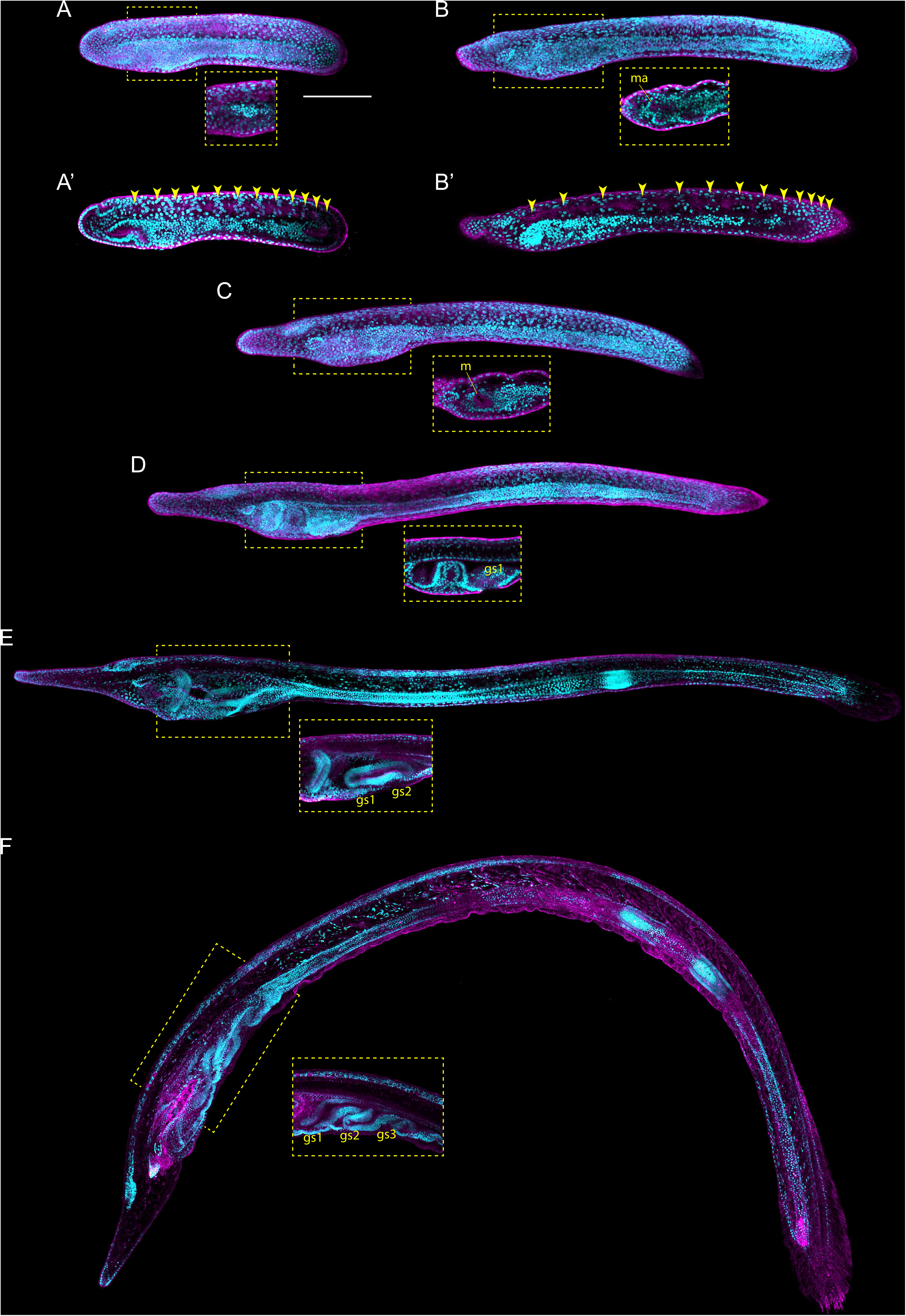
*Branchiostoma lanceolatum* tailbud and larval stages. Embryos and larvae are labeled for aPKC (magenta) and stained with the DNA dye Hoechst (cyan). Average projections for aPKC (magenta) and maximum projections for Hoechst DNA staining (cyan) of confocal z-stacks of entire embryos and larvae. Anterior pole is to the left and dorsal side is up. (A,A’) T0 stage, (B,B’) T1 stage, (C) L0 stage, (D) L1 stage, (E) L2 stage, (F) L3 stage. (A’,B’) Single z-stacks highlighting the inner morphology of the developing tailbud. Insets in (A-F) highlight the pharyngeal region. In (A’,B’), yellow arrowheads indicate the posterior limits of the somites. Abbreviations: gs1 – 1^st^ gill slit; gs2 – 2^nd^ gill slit; gs3 – 3^rd^ gill slit; m – mouth; ma – mouth anlagen. Scale bar: 100 μm.

The earliest larva, the L0 stage, already features the main structural elements that define the asymmetry, along the left-right axis, of all subsequent larval stages (Fig. 4C). The larval mouth opens on the left side of the developing animal by fusion of ectoderm and endoderm (Fig. 4C,C’) (Kaji et al., 2016; Holland, 2018). The left diverticulum has now penetrated the ectoderm to form the pre-oral pit, also known as Hatschek’s pit (Supplementary Fig. 2C). Hatschek’s nephridium, the kidney of larval lancelets, is now detectable between the ectoderm and the anterior-most somite on the left side of the larva (Hatschek, 1893; Holland, 2018). On the right side, the club-shaped gland is forming in the anterior endoderm, opposite to the mouth (Supplementary Fig. 2C) (Goodrich, 1930). Once completely developed, the club-shaped gland resembles a tube that connects the pharyngeal lumen on the right with the external environment on the left (Jefferies, 1987). The opening is located just anterior to the mouth and is characterized by cells bearing large cilia that create a water current from the exterior into the organ (Olsson, 1983). The club-shaped gland has been shown to secrete mucoproteins and might thus contribute to larval feeding (Holland, 2015). Another structure detectable on the right side of the pharynx at the L0 stage is the endostyle. The endostyle forms from a thickening of the endodermal wall and is located just anterior to the club-shaped gland (Supplementary Fig. 2C). The endostyle, which secretes mucous used to trap food particles, has been proposed to be homologous to the vertebrate thyroid gland (Ogasawara, 2000; Paris et al., 2008; Bertrand and Escrivá, 2011).

Although the definitive gill slits of lancelet larvae are found on the right side of the body (Holland, 2015), the anlage of the first gill slit forms at the ventral midline at the L0 stage (Supplementary Fig. 2C). The anlage of the anus arises at the same stage at the posterior end of the gut, which is located just anterior to the ectodermal caudal fin (Supplementary Fig. 2C) (Jefferies, 1987). However, while the anlage of the anus also originates at the ventral midline, the definitive anus will be located on the left side of the body (Jefferies, 1987). The first definitive gill slit penetrates at the L1 stage (Fig. 4D), and, following the establishment of all the structures referred to above, the L1 larva starts feeding. Following the L1 stage, new gill slits are added sequentially, hence defining the subsequent developmental stages: L2 stage for 2 gill slits (Fig. 4E), L3 stage for 3 gill slits (Fig. 4F) and so on, until the larva enters metamorphosis. The number of gill slits required before a larva becomes competent to undergo metamorphosis varies between different lancelet species (Wickstead, 1967; Holland and Yu, 2004; Urata et al., 2007; Carvalho et al., 2017b).

### 3.2. *Branchiostoma lanceolatum* developmental timing

It is well established that temperature directly affects the speed and potentially even the progression of animal development, in lancelets as well as in other animals (Fuentes et al., 2007; Ebisuya and Briscoe, 2018). To define the impact of temperature on *B. lanceolatum* development, we reared embryos and larvae at three different temperatures (16°C, 19°C and 22°C). We then mapped their developmental progression, according to our staging system and using somite pairs as defining landmark. To visualize the somites, embryos were fixed at regular intervals starting at the N0 stage, and *in situ* hybridization was performed with the somite marker *mrf1*. For each of the three temperatures, the number of somite pairs at a given developmental time was subsequently used as a training set (Supplementary Fig. 3, Supplementary Table 1) to define the growth curve that best reflected *B. lanceolatum* development. We further extrapolated the time intervals for the different development stages of our staging system prior to and following the neurula stages (Fig. 5). The results show that, despite a marked effect on the speed of development, the shapes of the growth curves, marking the progression of development, are very similar for the three temperatures (Fig. 5). This indicates that the different temperatures predominantly impact the rate of cell division during development and not the overall physiology of the embryos and larvae. It is, however, almost certain that *B. lanceolatum* can only develop within a certain temperature range. *B. lanceolatum* adults, for example, die after being cultured at 30°C for two weeks (Fuentes et al., 2007), and it is likely that embryos and larvae are even more temperature sensitive than adults. The results further demonstrate that these growth curves can be used to easily transform a developmental stage expressed as time after fertilization into an unambiguous stage name.

**Figure 5.**
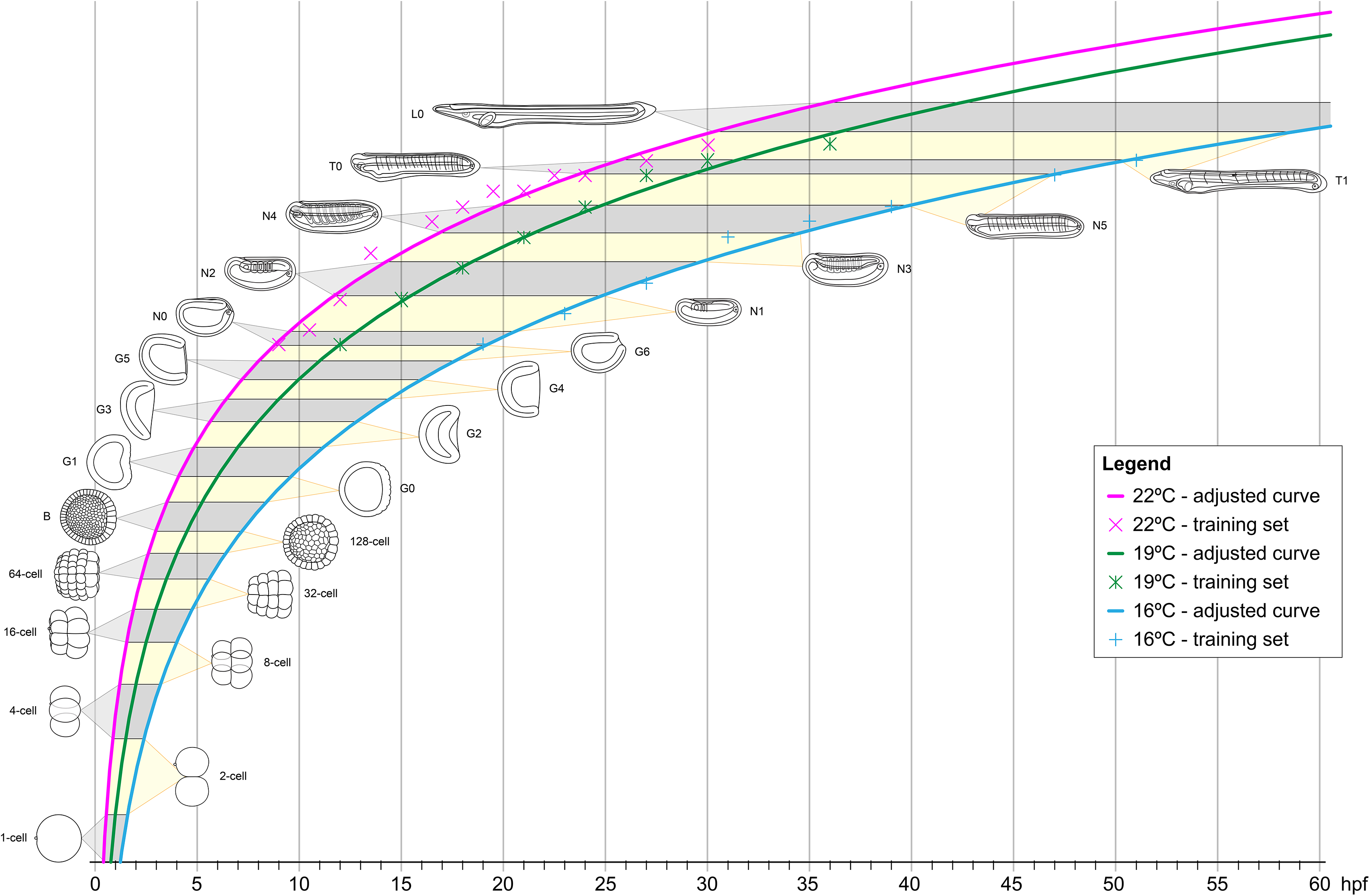
Growth curves of *Branchiostoma lanceolatum* embryos and larvae at 16°C, 19°C and 22°C. Animal schematics and stage nomenclatures are according to the staging system detailed in Figure 6. Tendency adjusted curves were obtained from the training sets and are defined by the equations: [y=12.403ln(x) - 36.493] for 16°C; [y=12.466ln(x) - 30.812] for 19°C; [y=11.25ln(x) - 24.354] for 22°C. These curves use natural logarithms and do thus not reach 0 hours post fertilization (0 hpf). The graphs were simplified accordingly. Abbreviations: hpf – hours post fertilization.

### 3.3. Comparative lancelet developmental staging

We next assessed whether the staging table we elaborated using *B. lanceolatum* (Fig. 6) can be applied to the development of other lancelets. For this, we compared *B. lanceolatum* embryos and larvae with those from four additional lancelet species, three from the genus *Branchiostoma* (*B. floridae*, *B. belcheri*, *B. japonicum*) and one from the genus *Asymmetron* (*A. lucayanum*). A total of 13 developmental stages were included in the comparative analysis: unfertilized eggs, 8-cell, 64-cell, 128-cell, B, G1, G4, G6, N1, N2, N4, T1 and L2 (Fig. 7). DIC images of the different stages revealed a strong overall conservation of the morphology of the five species. However, differences were detected in the overall size of the developing lancelets. The unfertilized egg of *B. floridae*, for example, is significantly larger than those of the other analyzed species. The diameter of the *B. floridae* egg is about 25% larger than that of *B. lanceolatum*, 18% larger than that of *B. belcheri*, 22% larger than that of *B. japonicum* and 33% larger than that of *A. lucayanum* (Fig. 7A). Another notable difference is the appearance of pigmentation in the posterior-most ectoderm, which is detectable as early as the N4 stage in *A. lucayanum*, but only appears at the T1 stage in the *Branchiostoma* species (Fig. 7B,C). In addition, the timing of rostrum and tail fin formation is not strictly conserved (Fig. 7C). Thus, while the rostrum is clearly elongated in T1 stage *B. lanceolatum*, development of the snout region is much less advanced in the other species, in particular in *A. lucayanum* (Fig. 7C). The lack of anterior head cavities in members of the genus *Asymmetron* may at least partially explain this prominent difference (Holland and Holland, 2010; Holland et al., 2015). Posteriorly, pigmented cells are detectable in *A. lucayanum* as well as *B. lanceolatum* and *B. belcheri*. In these three lancelet species, the rudiment of the forming tail fin is also already present at the T1 stage (Fig. 7C). In the larva, the species-specific differences in the snout and tail regions become even more accentuated. While *B. lanceolatum* larvae have a particularly long and thin snout, the rostrum of the other lancelet species is much less pronounced. At the L2 stage, the tail fins are either pointy (in *lucayanum*, *B. lanceolatum* and *B. belcheri*) or roundish (in *B. floridae* and *B. japonicum*). Previous studies have further shown that, when compared to *B. floridae*, *B. lanceolatum* larvae are characterized by a heterochronic delay of second gill slit formation and that this delay is not due to differences in developmental speed (Somorjai et al., 2008).

**Figure 6.**
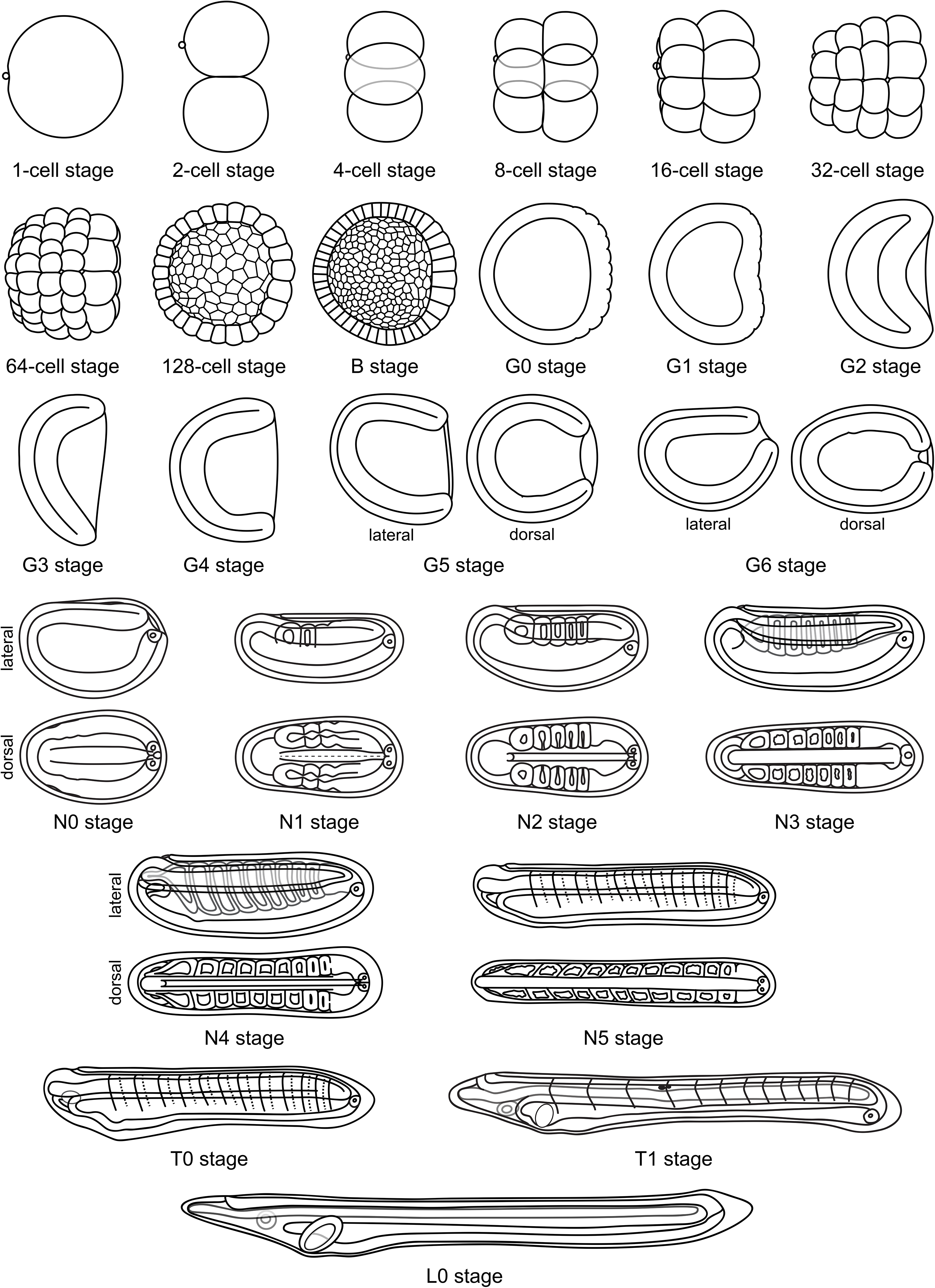
Schematic representation of *Branchiostoma lanceolatum* development. Representations from the 1-cell stage to the L0 stage. Animal pole and anterior pole are to the left and dorsal side is up in lateral views. Drawings adapted from Hatscheck’s original descriptions of *Branchiostoma lanceolatum* development (Hatschek, 1881).

**Figure 7.**
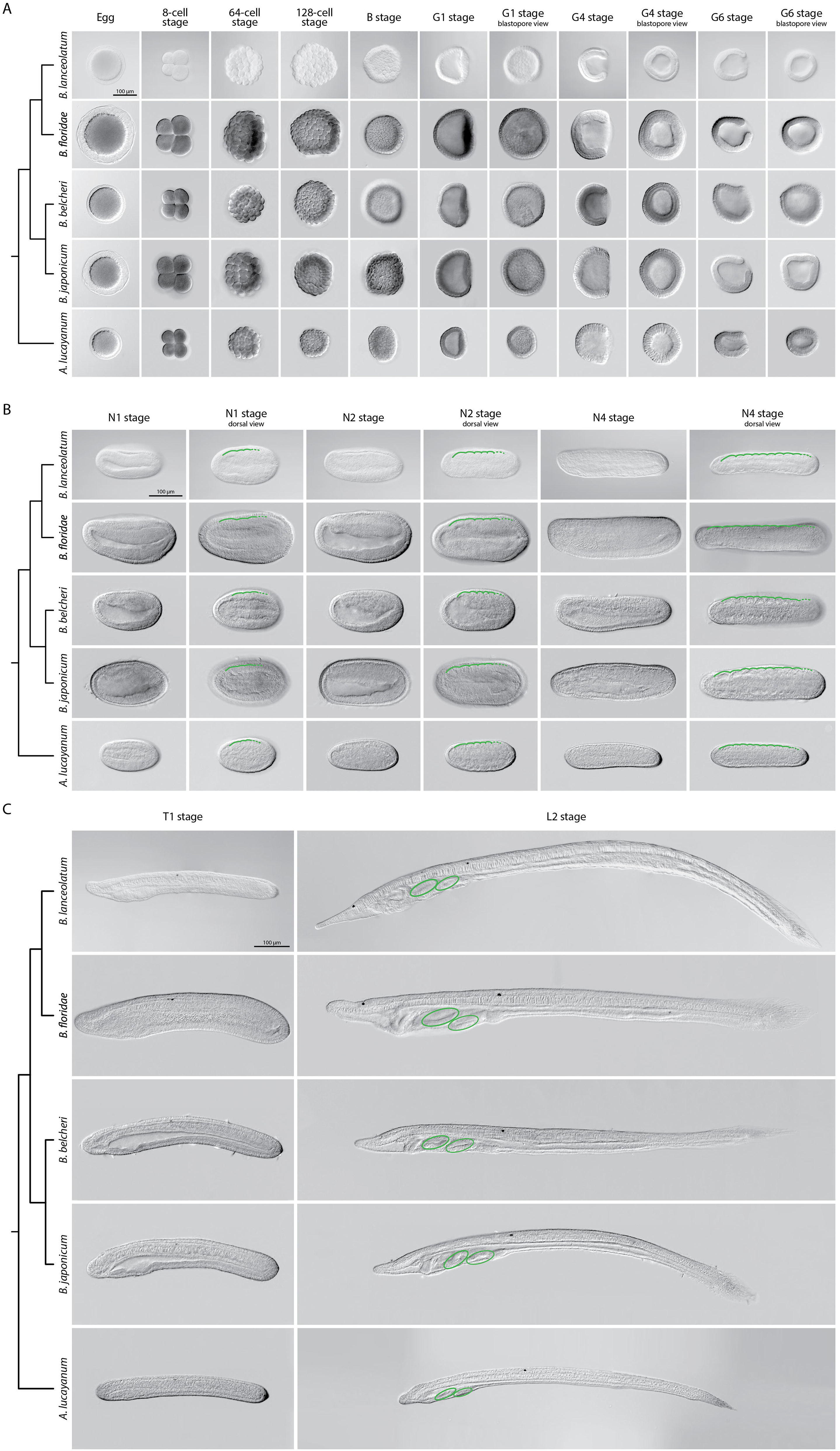
Comparison of lancelet development. Five species were analyzed: *Branchiostoma lanceolatum*, *Branchiostoma floridae*, *Branchiostoma belcheri*, *Branchiostoma japonicum* and *Asymmetron lucayanum*. (A) cleavage, blastula and gastrula stages, (B) neurula stages, (C) tailbud and larva stages. Cladograms represent the evolutionary relationship between the different species (Igawa et al., 2017). The green lines in (B) trace the somites on one side of the neurula, with dashed green lines highlighting forming somites. The green ovals in (C) indicate the gill slits of the larva. Scale bars: 100 μm.

Despite these differences, the defining characters of each developmental stage that we established in *B. lanceolatum* embryos and larva were conserved in all other lancelet species. The cleavage, gastrula and neurula stages of the five lancelet species are thus remarkably similar (Fig. 7A,B). Furthermore, the rate of somite formation as well as the timing of appearance of key morphological features at the neurula and tailbud stages are comparable (Fig. 7B,C). For example, the N2 stage embryo of all five species is characterized by 4 to 5 somite pairs, a neuropore and a neurenteric canal. Taken together, although there are minor species-specific differences, the overall development of the five lancelets is highly conserved and fully compatible with our updated staging and stage nomenclature systems. We thus expect these systems to be widely applicable to embryos and larvae of all extant lancelets.

## 4. Discussion

In the present study, we carried out a detailed analysis of the development of the lancelet *B. lanceolatum* using confocal microscopy and we defined straightforward staging and nomenclature systems for developing lancelets. We validated the updated staging system at different rearing temperatures for *B. lanceolatum* and demonstrated that it can be used for staging lancelets from the genus *Branchiostoma* as well as from the genus *Asymmetron*. This work thus resolves two fundamental problems for studies carried out in lancelets: (1) the lack of comparability between embryos and larvae from different species and (2) the confusion created by varying staging and stage nomenclature systems in a given species. Importantly, the morphological characters used to define each stage are generally easy to identify, such as the total number of cells for the cleavage stages, the initiation of asynchronous cell division for the blastula (B) stage, the shape of the gastrula (G), the number of somite pairs in the neurula (N) and tailbud (T) stages and the formation of pharyngeal structures for the tailbud (T) and larva (L) stages. Most of these characters have previously been validated as distinguishing hallmarks of lancelet development (Kovalevsky, 1867; Hatschek, 1893; Cerfontaine, 1906; Conklin, 1932; Hirakow and Kajita, 1990, 1991, 1994) and are also regularly used for the staging of other model organisms (Kimmel et al., 1995; Richardson and Wright, 2003).

Our updated staging system also allowed us to clarify previously unresolved controversies about lancelet development. One example is the definition of the blastula stage. Some authors suggested that the blastula is established as soon as the blastocoel is enclosed by cells (at the 64-cell stage) (Holland and Yu, 2004), while others proposed that the blastula forms after the 8^th^ round of cell divisions (after the 128-cell stage) (Hirakow and Kajita, 1990). Here, we redefined the B stage, which is characterized by the initiation of asynchronous cell divisions (at the transition from 128 cells to 256 cells) and ends with the initial flattening of the vegetal side of the embryo. In chordates, the first asynchronous cell divisions are often observed around the mid-blastula transition (MBT) and are thus correlated with the activation of zygotic gene transcription (McDougall et al., 2019). A detailed analysis of transcriptomes obtained at different developmental stages suggests that this is also the case in amphioxus, as the transition from 128 cells to 256 cells is marked by a strong increase in the expression of genes required for the initiation of zygotic transcription, including, for example, those encoding nuclear ribonucleic proteins (Yang et al., 2016).

Another ambiguous developmental period is the transition between the gastrula and the neurula stage, sometimes referred to as a very late gastrula (Hirakow and Kajita, 1991) or a very early neurula (Lu et al., 2012; Zhang et al., 2013). We redefined this important stage as N0, corresponding to an embryo with a small blastopore, which is characteristic for gastrula stages, and a flattened neural plate, marking the onset of neurulation. We further expanded the classification of neurulae to six independent N stages, hence allowing more detailed descriptions of the morphological changes occurring during this crucial developmental period. Previous descriptions distinguished only three (Hirakow and Kajita, 1994) or four different N stages (Lu et al., 2012).

Another controversial point of lancelet development is the definition of the larva. Some authors claimed that the larval stage starts when “tissues and cells prepare for performing their own function” (Hirakow and Kajita, 1994). Alternatively, the larval stage has been defined by the opening of the mouth and thus by the moment the animal starts feeding (Holland, 2015). To clarify this issue, we defined a new developmental period for lancelets that, based on the gestalt of the embryo at this stage, we called the tailbud (T) stage (Lemaire, 2011). We further defined the onset of the larval stage (L0) as the moment when the mouth opens, as it has previously been suggested for lancelets (Holland, 2015) and other animals (Kimmel et al., 1995; Smith et al., 2008).

Significant efforts have been made to develop protocols for maintaining and spawning adult lancelets in captivity and for manipulating lancelet embryos and larvae. Thanks to these efforts, lancelets have become attractive laboratory models (Carvalho et al., 2017b; Su et al., 2020). However, one of the remaining obstacles was the absence of a widely applicable staging system guaranteeing the comparability of results obtained in different lancelet species. Here, we propose a complete staging system for developing lancelets. Although the stage descriptions were carried out in *B. lanceolatum*, our comparisons with other lancelet species clearly demonstrate that both staging and nomenclature are valid beyond *B. lanceolatum* and are likely applicable to all extant lancelets. Using the defining characters for each stage, we were thus able to establish a comparative developmental table for the five lancelet species used in this study: *B. lanceolatum*, *floridae*, *B. belcheri*, *B. japonicum* and *A. lucayanum* (Table 1). In this regard, this work adds morphological evidence to genomic results suggesting that lancelets evolve at a very slow rate (Putnam et al., 2008; Igawa et al., 2017; Marlétaz et al., 2018; Simakov et al., 2020). Taken together, we strongly believe that this description and organization of embryonic and larval development, along with the ontology for anatomy and development for the *Branchiostoma* genus (AMPHX) (Bertrand et al., 2021), should become the standards for the scientific community in an effort to harmonize research on developing lancelets. We also anticipate that this updated description of lancelet development will facilitate future comparative studies between lancelets and other chordates.

**Table 1.** Comparison of lancelet development. Species: *Branchiostoma lanceolatum*, *Branchiostoma floridae*, *Branchiostoma belcheri*, *Branchiostoma japonicum*, *Asymmetron lucayanum*. Data origin: ^1^ Current study, ^2^ Stokes and Holland, 1995; Holland and Holland, 1998; Holland and Yu, 2004; Holland et al., 2015, ^3^ Zhang, 2017, ^4^ Hirakow and Kajita, 1990, 1991, 1994; Morov et al., 2016, ^5^ Holland and Holland, 2010; Holland et al., 2015. “/” indicates that different developmental times have been reported.

|  |  | *B. lanceolatum*<sup>1</sup> |  |  | *B. floridae*<sup>2</sup> |  | *B. belcheri*<sup>3</sup> | *B. japonicum*<sup>4</sup> | *A. Lucayanum*<sup>5</sup> |
| --- | --- | --- | --- | --- | --- | --- | --- | --- | --- |
| Stage | Key feature | at 16°C | at 19°C | at 22°C | at 25°C | at 30°C | at 24°C | at 25°C | at 27°C |
| 1-cell stage | fertilized egg |  |  |  |  |  |  |  |  |
| 2-cell stage | 2 cells |  | 1h |  | 45' | 30' | 45' | 50'-1h | 1h-1h30' |
| 4-cell stage | 4 cells |  | 1h30' |  | 1h | 50' | 1h | 1h10' | 2h-2h30' |
| 8-cell stage | 8 cells |  | 2h |  | 1h30' | 1h | 1h20' | 1h35' | 2h-2h30' |
| 16-cell stage | 16 cells |  | 2h30' |  | 2h | 1h15' | 1h45' | 1h55' | 3h-3h30' |
| 32-cell stage | 32 cells |  | 3h |  | 2h15' | 1h30' | 2h10' | 2h15' | 3h-3h30' |
| 64-cell stage | 64 cells |  | 3h30' |  | 2h30' | 1h45' | 2h30' | 2h35' | 4h-4h30' |
| 128-cell stage | 128 cells |  | 4h |  | 3h | 2h | 2h50' | 3h10' | 4h-4h30' |
| B stage | initiation of asynchronous cell division |  | 4h30' |  | 4h | 2h30' |  | 3h20' | 5h |
| G0 stage | initial flattening of the vegetal zone |  | 5h |  |  |  | 3h30' |  |  |
| G1 stage | flattened vegetal pole |  | 6h |  | 4h30' | 3h30' |  | 3h40' |  |
| G2 stage | invaginated vegetal pole |  | 7h |  |  |  |  | 4h10' |  |
| G3 stage | cap shaped |  | 8h |  | 5h | 4h | 3h55' | 5h35' |  |
| G4 stage | cup shaped | 15h | 9h |  | 6h | 4h30' |  | 6h-6h20' | 9h |
| G5 stage | vase shaped |  | 10h |  |  |  | 5h45' | 7h40' |  |
| G6 stage | bottle shaped |  | 11h |  | 6h30' | 5h | 7h55' | 8h50' |  |
| N0 stage | no somite pairs, neural plate | 19h | 12h | 9h |  |  | 8h30' |  | 12h |
| N1 stage | 1-3 somite pairs | 23h | 15h | 12h | 8h30' | 6h | 10h20' | 10h30' | 15h |
| N2 stage | 4-5 somite pairs, hatching | 27h | 18h |  | 9h30' | 6h30' | 12h | 13h | 19h |
| N3 stage | 6-7 somite pairs | 31h | 21h | 13h30' |  |  |  |  |  |
| N4 stage | 8-9 somite pairs,<br>prior to schizocoelic somite formation | 35h | 24h | 16h30' | 10h30' | 7h30' |  | 18h |  |
| N5 stage | 10-11 somite pairs | 47h | 27h | 19h30' |  |  |  |  | 32h |
| T0 stage | 12 somite pairs, tailbud shape,<br>enlarged pharyngeal region | 51h | 30h | 27h |  |  |  |  |  |
| T1 stage | 13 somite pairs, mouth and pre-oral pit<br>anlagen, first pigment spot |  | 36h | 30h | 20h/24h | 12h/17h |  | 24h | 50h |
| L0 stage | no gill slits, open mouth |  | 42h |  | 30h | 21h |  |  |  |
| L1 stage | 1 gill slit |  | 48h |  | 32h/36h | 23h/28h |  | 36h | 72h |
| L2 stage | 2 gill slits |  |  |  | 42h/72h | 36h | 36h/48h | 48h |  |
| Ln stage | n gill slits |  |  |  |  |  |  |  |  |

## Acknowledgements

The authors are indebted to Linda Z. Holland and Nicholas D. Holland from the Scripps Institution of Oceanography, La Jolla, USA, for collecting *Asymmetron lucayanum* adults. Janet Chenevert from the Laboratoire de Biologie du Développement de Villefranche-sur-Mer, Villefranche-sur-Mer, France, kindly provided the FM 4-64 lipophilic dye and the primary antibody against aPKC as well as useful technical advice. We would like to thank Estelle Hirsinger from the Institut de Biologie Paris-Seine, Paris, France, for reading and commenting the manuscript. We are further grateful to Tzu-Kai Huang and the staff at the Marine Research Station of the Institute of Cellular and Organismic Biology for technical assistance. We also thank the Centre de Ressources Biologiques (CRB) of the Institut de la Mer de Villefranche (IMEV), specifically the Service Aquariologie (SA) and the Mediterranean Culture Collection of Villefranche (MCCV), and the Plateforme d’Imagerie par Microscopie (PIM) of the Institut de la Mer de Villefranche (IMEV), which are supported by EMBRC-France (ANR-10-INBS-02). JEC was a FCT doctoral fellow (SFRH/BD/86878/2012) and is currently supported by a FRM fellowship (SPF20170938703). LWY and JKY are supported by Academia Sinica intramural funds and grants from the Ministry of Science and Technology, Taiwan (MOST-105-2628-B-001-003-MY3 and MOST-108-2311-B-001-035-MY3). HE is financed by ANR-16-CE12-0008-01, ANR-17-CE13-0027-02 and ANR-19-CE13-0011. MS is funded by the CNRS.

---

**Supplementary Figure 1** – Detailed highlights of specific structures of *Branchiostoma lanceolatum* development during cleavage and neurula stages. (A,B) Embryos are stained with the lipophilic dye FM 4-64 (magenta). (C-E) Embryos are labeled for aPKC (magenta) and stained with the DNA dye Hoechst (cyan). The anterior pole is to the left, and, on the dorsal views, the right side is up (C,C’,E-E’’), while, on the lateral view, the dorsal side is up (D). Maximum projections of confocal z-stacks of *B. lanceolatum* embryos at the 1 cell-stage (A), 32-cell stage (B), N1 stage (C,C’) and N2 stage (D-E’’). Insets correspond to regions highlighted with dotted rectangles and are shown at 2x magnification. Abbreviations: bc – blastocoel; bp – blastopore; cv – cerebral vesicle; m – maternal DNA; nc – neurenteric canal; np – neuropore; nrt – neural tube; nt – notochord; p – paternal DNA; pb1 – 1^st^ polar body; pb2 – 2^nd^ polar body; phc – presumptive head cavities; s1-5 – somite pairs 1 to 5; sm – somitic mesoderm. Scale bar: 50 μm.

**Supplementary Figure 2** – Detailed highlights of specific structures of *Branchiostoma lanceolatum* development during tailbud and larval stages. Embryos and larvae are labeled for aPKC (magenta) and stained with the DNA dye Hoechst (cyan). Embryos and larvae are in lateral views, the anterior pole is to the left and the dorsal side is up. T0 (A), T1(B) and L0 (C) stages are shown. Insets correspond to regions highlighted with dotted rectangles and are shown at 2x magnification. Abbreviations: an – anus; cc – cephalic coelom; csg – club-shaped gland; cv – cerebral vesicle; en – endostyle; np – neuropore; nrt – neural tube; nt – notochord; pgs – presumptive 1^st^ gill slit; pp – pre-oral pit; ps – 1^st^ pigment spot; rd – right diverticulum; s2-5 – somite pairs 2 to 5; tf – tail fin. Scale bar: 50 μm.

**Supplementary Figure 3** – Expression of the *mrf1* gene in developing *Branchiostoma lanceolatum* reared at different temperatures. Embryos are in dorsal views with anterior pole to the left and right side up. (A) 16°C, (B) 19°C and (C) 21°C. On each image, the time of development in hours post fertilization (h) and the number of fully formed somite pairs (s) are indicated. Scale bars: 50 μm.

**Supplementary Table 1** – Somite pair counts based on the expression of the *mrf1* gene in developing *Branchiostoma lanceolatum* reared at three different temperatures (16°C, 19°C and 21°C) (Supplementary Fig. 3), and natural logarithmic tendency curves obtained from the three training sets.

