## Supplementary figures and images for "An updated staging system for cephalochordate development: one table suits them all"

### Supplementary Fig.1

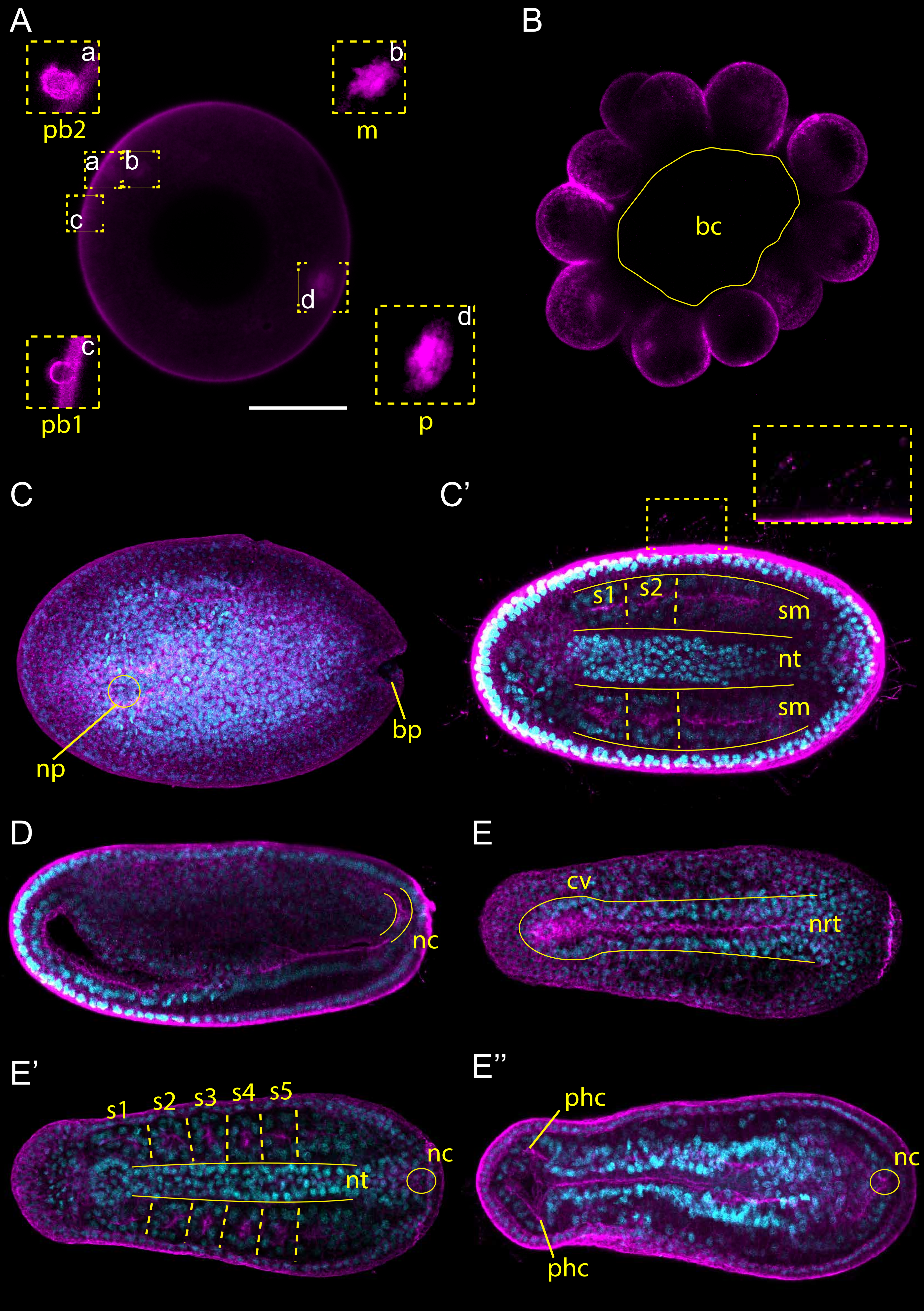

### Supplementary Fig. 2

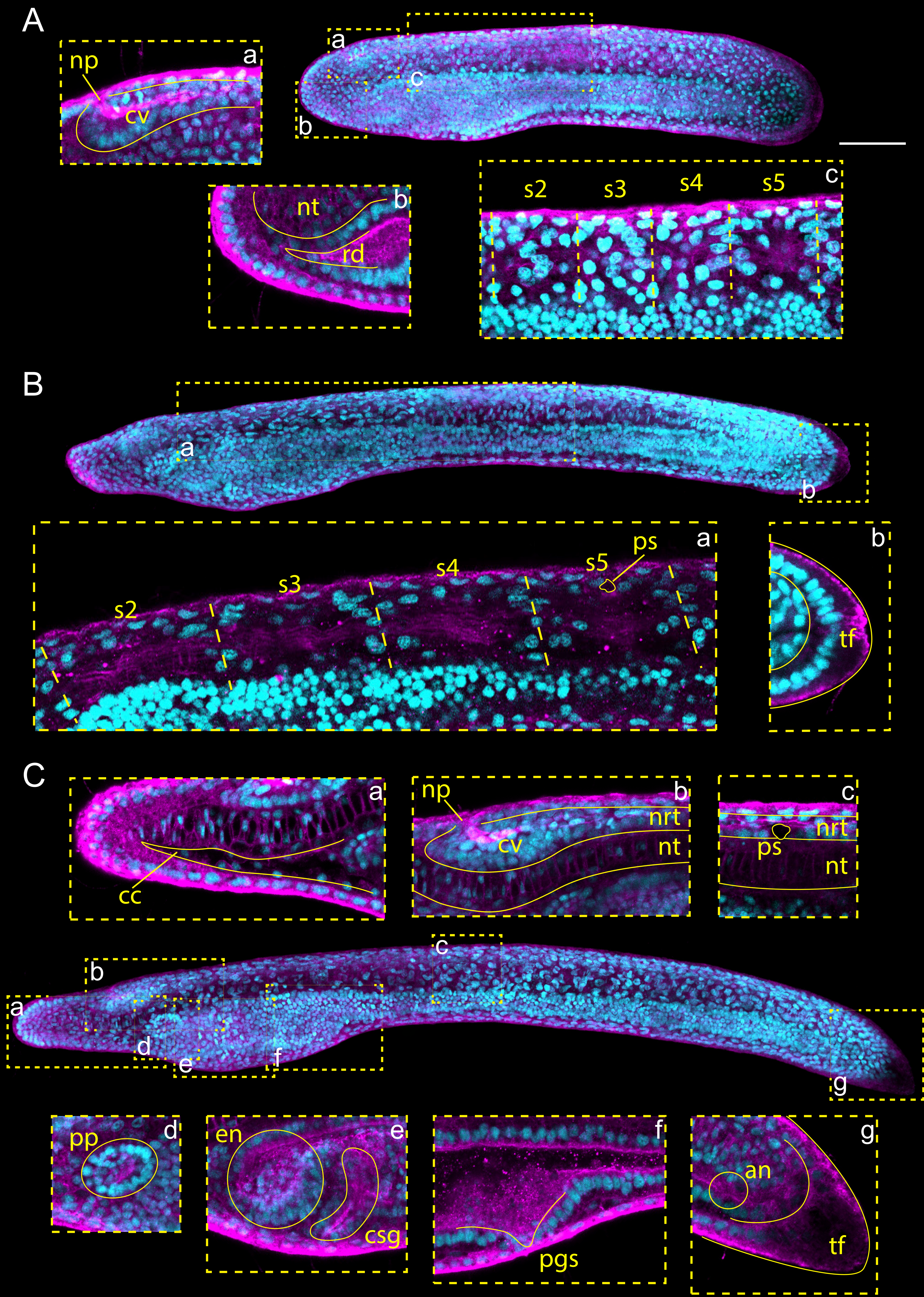

### Supplementary Fig. 3

Supplementary Figure 1

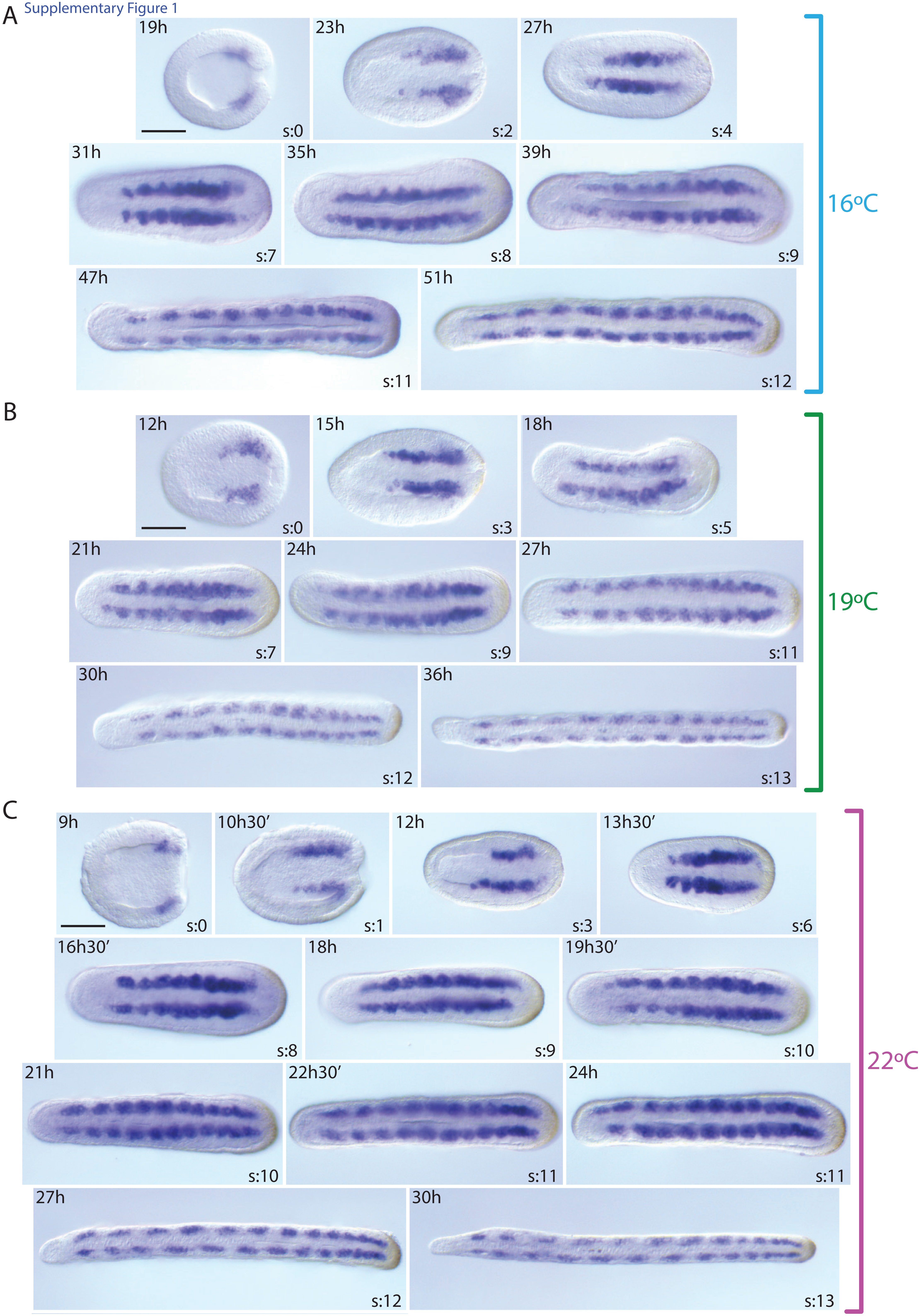
